# Noninvasive tomography of tumor progression via dynamical changes of T-cells in surrounding microenvironment

**DOI:** 10.64898/2026.09.19.752909

**Authors:** Longqiang Xu, Pinchao Meng, Weishi Yin, Hongyu Liu

## Abstract

Tumor progression is shaped by a continuous competition between malignant expansion and immune access within the surrounding microenvironment. Although spatial assays can reveal where T cells accumulate, most current approaches remain descriptive, treating immune organization as a static pattern and failing to infer the latent resistance structure that governs progression. Here we show that changes in peri-tumoral T-cell distribution can support noninvasive tomography of tumor progression. Using spatially resolved human breast tumor sections and controlled numerical experiments, we formulate immune-tumor interaction as a mean-field game (MFG) in which T cells act as agents responding to a heterogeneous microenvironment while the tumor shapes an equivalent barrier field that restricts infiltration. This framework converts boundary enrichment, layered blocking and core penetration patterns into a constrained inverse problem whose solution yields quantitative maps of immune exclusion, barrier strength and progression-associated internal states. Forward simulations show that barrier geometry reshapes immune-density equilibria, and inverse experiments show that closely related barrier fields can be recovered from immune-density observations under controlled and tissue-shaped proxy settings. By linking observable T-cell redistribution to latent microenvironmental resistance, this approach establishes a route toward noninvasive, repeatable assessment of tumor progression without direct biopsy of the underlying barrier structure.

## 1 Introduction

Tumor progression depends on interactions between malignant tissue and the surrounding immune microenvironment. Stromal architecture, extracellular matrix, suppressive cues and tissue geometry influence whether T cells approach, traverse and enter malignant regions [1–3]. In breast cancer, both the abundance and localization of tumor-infiltrating lymphocytes carry biological and clinical information, motivating standardized separation of stromal and intratumoral compartments [4, 5]. T-cell abundance alone therefore provides an incomplete account of immune access; location relative to the tumor boundary, peri-tumoral layers and core is also informative.

Spatial transcriptomics and computational pathology pipelines quantify enrichment, distance, spatial organization and clustering [6–10]. These measurements describe where T cells accumulate; connecting them to a latent resistance representation requires an explicit population model that links spatial density to movement cost and collective interaction.

We reasoned that peri-tumoral T-cell redistribution should be interpreted as the outward signature of an underlying immune-tumour competition law. If the tumour and its surrounding microenvironment generate a heterogeneous resistance landscape, then the observed T-cell pattern should retain information about that hidden structure. We therefore model immune access as a mean-field game in which T cells act as interacting agents seeking favorable migration paths, while the tumour shapes an equivalent barrier field that resists infiltration. This gives rise to an inverse mean-field-game problem. Here, the forward dynamics generate spatial redistribution patterns, and the inverse problem estimates the corresponding resistance field from observed T-cell distributions. We note that recent mathematical work has studied inverse mean-field-game problems in which unknown potentials, coefficients or interactions are inferred from observed population states [11], which provides a theoretical foundation for the present approach.

The biological and mathematical strands therefore meet at a specific unresolved interface. Spatial transcriptomics and pathology studies provide increasingly detailed descriptions of cell composition and local interaction, while inverse MFG studies provide tools for inferring latent model structure from population observations. We connect these strands by defining the inferred field as an equivalent functional resistance representation and evaluating its computability and stability in controlled numerical and structured tissue-record settings.

In this study, we develop a framework grounded in mean-field game theory to examine how the spatial redistribution of T cells reflects the influence of the tumour microenvironment on immune access. We test whether the framework can translate observed T-cell organization into a comparable functional state of immune resistance across controlled numerical settings and tumour sections with structured immune–tumour organization. This framework provides a mechanistic route for connecting observable immune organization with latent tumour–microenvironment states and establishes a basis for future noninvasive assessment based on spatial T-cell information.

## 2 Results

### 2.1 Dynamic and layered T-cell redistribution tracks tumor progression states

We examined whether peri-tumoral T-cell distribution contains reproducible information about tumor progression beyond overall immune abundance. Across human breast tumor sections, T cells did not simply vary in number; they redistributed differently relative to the tumor boundary, surrounding stromal layers and tumor core. Some regions showed broader penetration across the boundary and deeper extension toward malignant tissue, whereas others showed strong boundary accumulation, stratified depletion across peri-tumoral layers or near-complete exclusion from inner tumor regions.

These differences are not well summarized by cell count alone. Regions with similar T-cell density can differ sharply in progression-relevant accessibility: one can show sustained movement toward the core, while another shows high peripheral density but rapid decay across the boundary. To capture this structure, we define three linked observation channels: boundary infiltration, layered infiltration and core penetration. Boundary infiltration measures immune accumulation near the tumor margin; layered infiltration measures how that signal decays or persists across successive peri-tumoral zones; and core penetration measures the extent of effective immune access within inner tumor regions.

The channel definitions are conceptually related to the earlier pathological distinction between stromal and intratumoral TILs [4, 5]. Here they serve as spatial-transcriptomic marker proxies for immune access on a common computational grid and are distinct from a clinical TIL score.

The recurrence of these layered redistribution patterns suggests that T-cell organization carries spatial information beyond abundance, including where immune advance stalls, decays across layers or reaches the tumor core. These features form the observation target for tomographic inference.

The audited project records provide 102 breast spatial sections, of which 99 marker-complete, full-transcriptome sections from compatible spatial platforms form the primary section-level cohort. A predeclared sensitivity subset retains 84 sections with at least 1,000 in-tissue spots and 2,000 detected genes. Figure 1 shows the implemented data-to-physics chain: original tissue context and measured immune-module spots are followed by observation and context fields, an equivalent functional resistance field is inferred, and the stationary MFG system is solved again to produce a same-section density residual. Original tissue images are shown without analytical overlays, while the registered HT397B1 image is repeated once with the measured source spots. Constructed input fields use index-preserving, edge-sharing hexagonal display lattices oriented to registered upper-origin pixel axes, whereas equation-native solution fields remain on the regular PDE solver grid with a neutral background. In the breast and colorectal examples, the observation and solver-derived spot fields are additionally registered to the complete pixel frame of the adjacent source image. Throughout, *B̂* denotes the model-derived equivalent functional resistance field.

**Figure 1:**
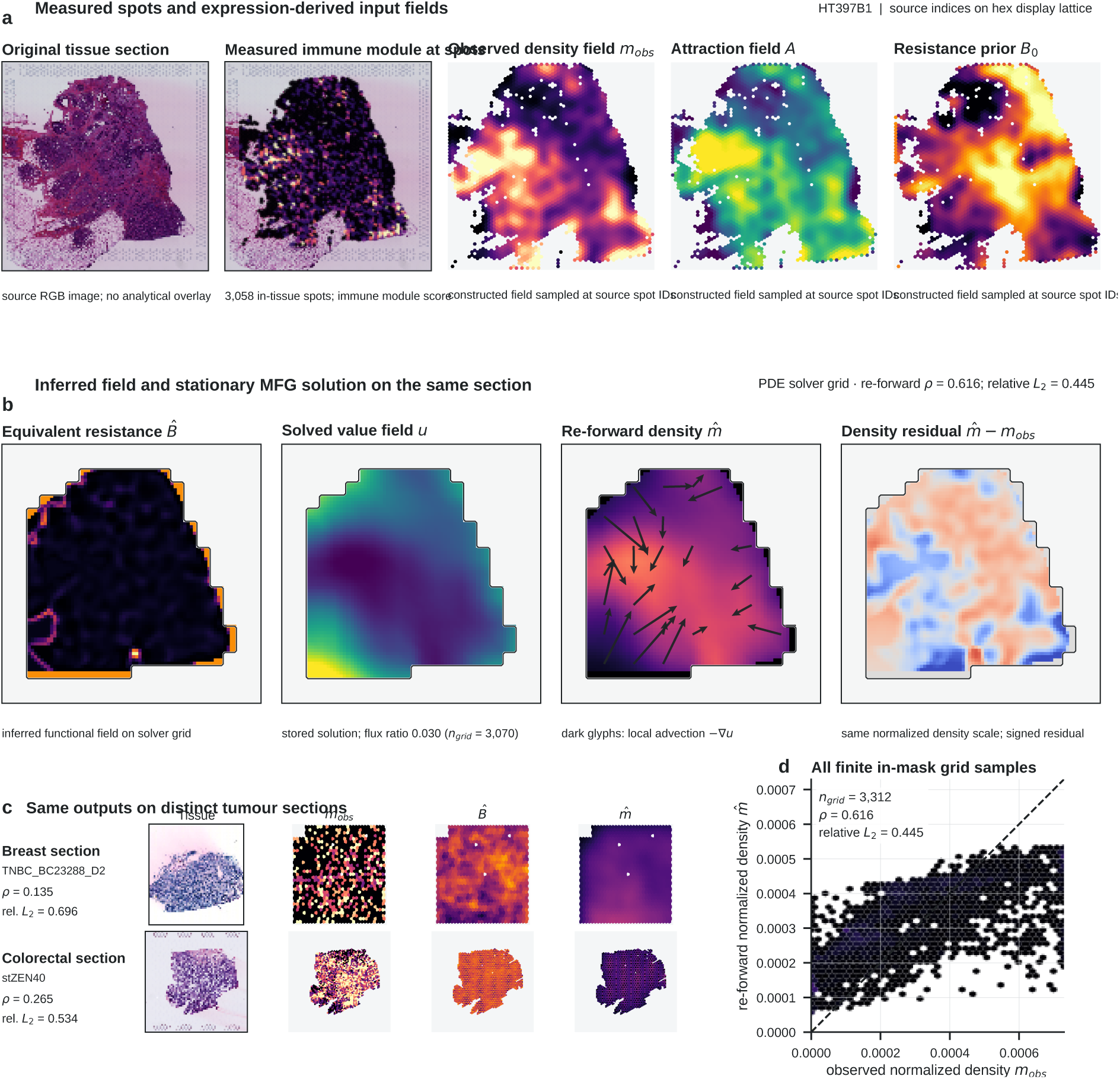
Breast spatial data and the MFG inverse workflow. a, Original HT397B1 tissue without an overlay, the same registered image with measured immune-module spots, and observed-density, attraction and resistance-prior inputs at 3,058 source spot IDs. Constructed fields use an index-preserving hexagonal display oriented to the tissue image. **b,** Equivalent resistance *B̂*, value field *u*, re-forward density *m̂* and signed residual on the regular PDE grid; dark glyphs show the calculated local *−∇u* direction. **c,** Original tissue images without overlays and the source observation, inferred field and re-forward output for breast sample TNBCBC23288D2 and colorectal sample stZEN40. Within each row, a registered-coordinate transform places the edge-sharing source-index tessellation in the same complete upper-origin image-pixel frame as the adjacent tissue image. **d,** Finite in-mask observed and re-forward density pairs with the identity line and section-level consistency metrics. Source data are provided.

### 2.2 Mean-field game dynamics convert T-cell redistribution into tumor tomography

We formulated observable redistribution as a mean-field-game inference problem in which T cells move through a heterogeneous tissue domain under the combined influence of migration cost, collective interaction and microenvironmental resistance. The tumor and its surrounding microenvironment are summarized by an equivalent barrier field that modulates the local difficulty of immune entry and propagation.

Toy steady-state numerical examples illustrate this logic directly: localized or multifocal barrier fields generate peripheral accumulation, layered depletion and reduced core access even before any inverse reconstruction is performed. These forward examples help distinguish the barrier field from a descriptive heatmap by showing how barrier geometry shapes the equilibrium immune distribution. Controlled forward simulations further showed that barrier topology and spatial layout reshape the immune-density equilibrium. Across five synthetic tumor configurations, distinct core, multifocal, lobulated, bridged and peripheral-arc barriers induced different redistribution patterns. In a real slice-derived proxy field, agreement between the forward MFG density and the observed T-cell proxy was sensitive to barrier strength: a weaker barrier setting improved the density–proxy correlation from 0.1271 at baseline to 0.2781, whereas a stronger barrier reduced it to 0.0203. These results support the use of the MFG system as a structured forward map from latent resistance geometry to observable immune organization, while also indicating that parameter calibration is important.

Given a candidate barrier field, the forward model predicts how T cells distribute across the boundary region, peri-tumoral layers and tumor core. The inverse problem identifies the field and associated parameters whose predicted patterns best match the observed multichannel data, thereby representing T-cell density as the outcome of constrained propagation under a competition law.

The resulting tomographic representation includes spatial resistance maps, blocking intensity across layers and summary parameters for modeled resistance to immune advance. These outputs can be compared across samples and regions under a common model specification.

### 2.3 MFG-constrained inverse tomography recovers barrier structure

We tested recovery under a standardized controlled benchmark in which five related single-peak resistance fields generated forward density observations and were then reconstructed with the stabilized MFG-constrained inverse procedure. Across all five predefined cases, correlations between the reference and recovered fields ranged from 0.9962 to 0.9990, while relative re-forward density errors ranged from 0.0833 to 0.1159 (Fig. 2a–c).

**Figure 2:**
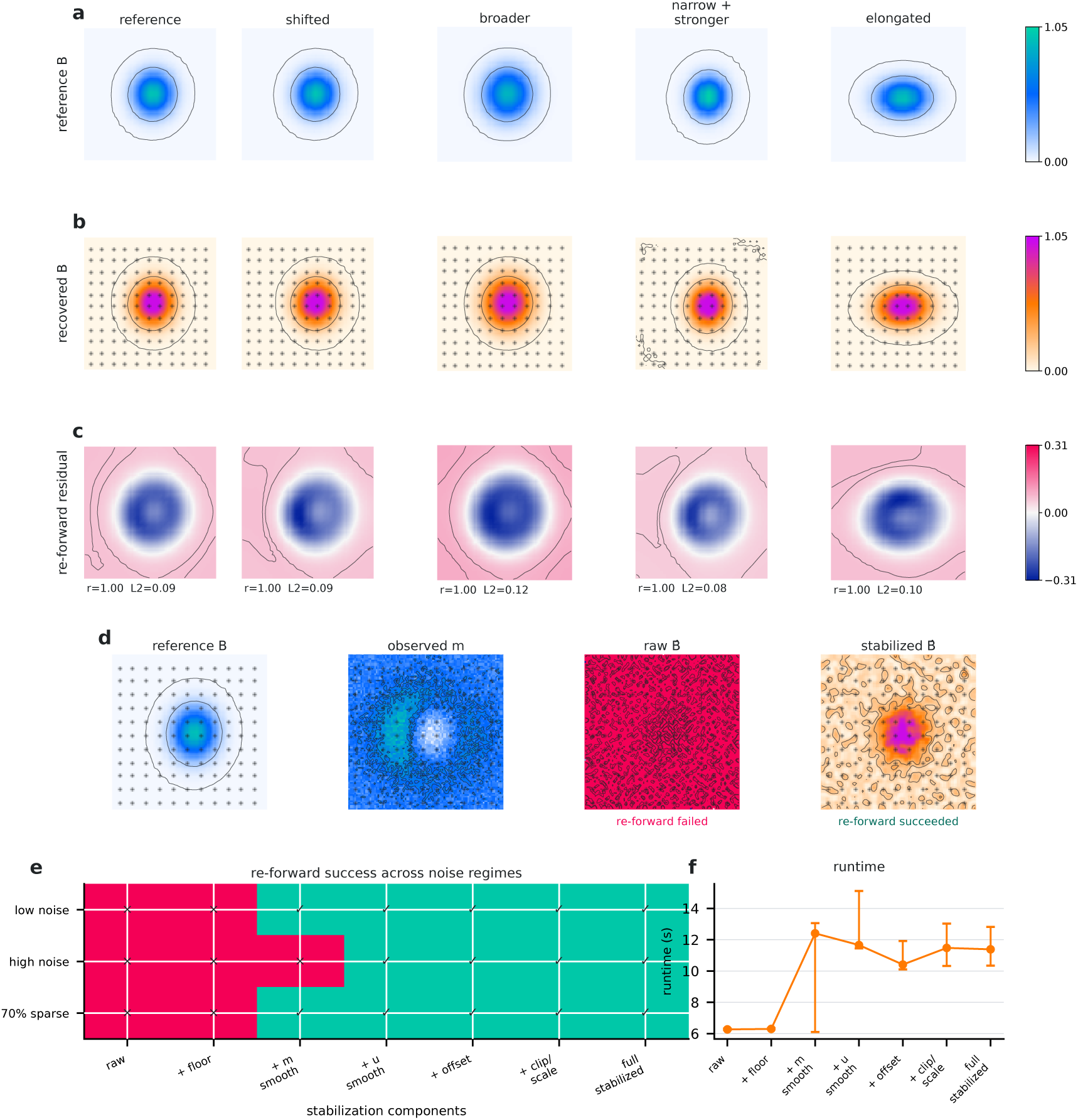
Stabilization preserves controlled resistance geometry and rescues degraded re-forward solves. a, Reference resistance fields for five predefined standardized similar-scene cases, with fixed field-quantile contours. **b,** Recovered equivalent fields *B* on the same cases, with sampled lattice points overlaid. **c,** Signed re-forward density residuals on one common symmetric scale; labels report the stored resistance correlation and relative re-forward *L*_2_ error for each case. **d,** The predefined high-noise case shown as reference *B*, observed density, raw *B* and stabilized *B*; the raw result fails re-forward solution and is clipped only for display, whereas the stabilized result succeeds. **e,** Complete 3-by-7 ordered stabilization lattice across the three degraded observations; crosses and circles denote failed and successful states. **f,** Runtime median and full minimum– maximum range across the three deterministic cases. All panels summarize deterministic numerical test cases. Source data are provided.

The shared-scale field plate shows that the stabilized inverse solutions retain the predefined differences in peak position, width, amplitude and elongation. Signed re-forward residuals are displayed on one common symmetric range, so spatial error patterns can be compared without panel-specific rescaling.

We also evaluated the numerical stabilization path under three predefined degraded observations. Raw and floor-only reconstructions failed re-forward solution in all three cases; density smoothing rescued two cases, and the addition of value-field smoothing was the first ordered variant to succeed in all three. In the predefined high-noise example, the raw recovered field reached a maximum of 14.9 and failed re-forward solution, whereas the stabilized result succeeded with resistance-field correlation *r* = 0.941 and relative re-forward *L*_2_ error 0.053 (Fig. 2d). These three deterministic numerical cases isolate the effects of the stabilization sequence.

### 2.4 Breast-section re-forward fit is heterogeneous across sections and sources

We assessed numerical re-forward consistency in the complete 99-section primary breast cohort. Stabilized re-forward fields were available for 96 sections, while three external-source sections failed at the forward-prior stage and remained in the cohort accounting. Among the 96 successful sections, the section-level correlation between observed and re-forward density fields had a median of *−*0.060 (interquartile range, *−*0.115 to 0.046; range, *−*0.380 to 0.428; Fig. 3a). Cohort-wide re-forward consistency was therefore weak under the current parameterization.

**Figure 3:**
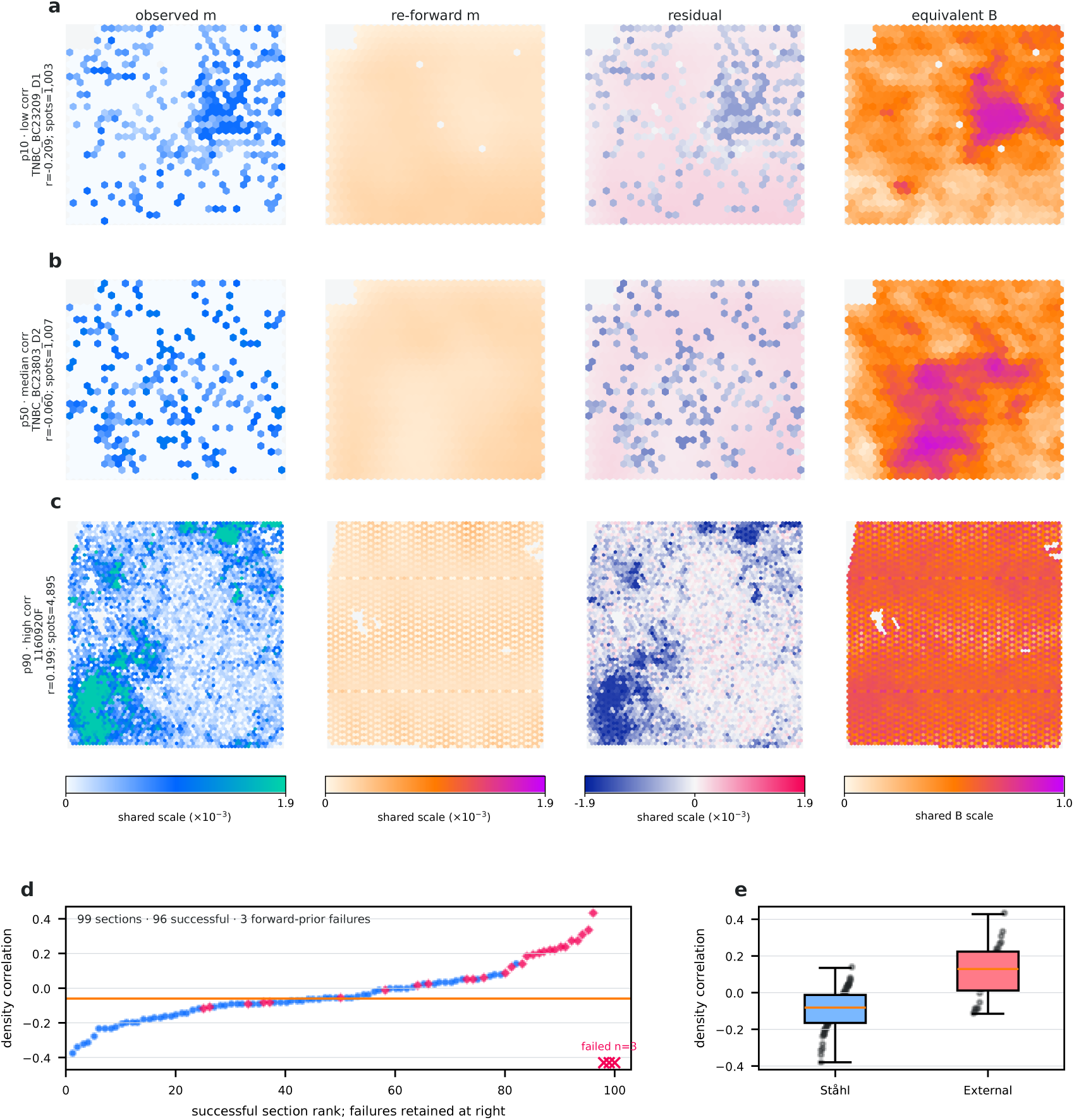
Breast-section re-forward fit is heterogeneous across sections and data sources. a–c, Successful breast sections at the p10, p50 and p90 ranks of density correlation. Each row shows direct source-spot observed density, re-forward density sampled at the same IDs, pointwise signed residual and sampled equivalent resistance *B* on a pointy-top, edge-sharing hexagonal display lattice. Stored array indices are preserved and missing indices remain blank. Observed and reforward density share a 0–0.0019 range, residuals share a symmetric *−*0.0019–0.0019 range and all three *B̂* fields share a 0–1.0 range. **d,** Ranked density correlations for all 96 successful sections among the 99-section cohort; red crosses denote the three forward-prior failures without assigned fit values. **e,** Source-stratified section-level distributions with individual sections shown. Spatial section is the statistical unit; source comparisons are descriptive. Source data are provided.

The predeclared sensitivity cohort contained 84 sections, of which 82 produced successful re-forward fields; its median density correlation was *−*0.058 (Fig. 3b). Source-stratified distributions were also heterogeneous: all 68 Ståhl sections succeeded, whereas 28 of 31 external-source sections succeeded and three failed (Fig. 3c). These source summaries are descriptive because a between-source hypothesis test was not predefined and complete verified patient grouping is unavailable for the external sections.

Successful sections were ranked by density correlation, and those nearest the p10, p50 and p90 ranks were selected for spatial inspection. Their direct source-spot observations, solver-derived fields sampled at the same spot IDs, residuals and equivalent fields are shown using shared data-derived ranges without per-panel normalization (Fig. 3a–c). All three rows use a pointy-top, edge-sharing hexagonal display lattice mapped from stored array indices; the layout preserves index adjacency for display, while absent source indices remain blank. The cohort rank strip and source-stratified section-level distributions are shown in Fig. 3d,e.

### 2.5 Attraction structure limits identifiability of the equivalent resistance field

To assess separation of resistance from the specified attraction structure, we compared *B* with the attraction field and barrier prior across all 99 primary breast sections. The median correlation was 0.825 for attraction and 0.454 for the barrier prior. Under the predefined descriptive rules, 60 sections were labelled severe A/B confounding, 38 attraction-dominant ambiguous and one moderate A/B confounding (Fig. 4a,b). Thus, *B* remained strongly coupled to the specified attraction field under the current model.

**Figure 4:**
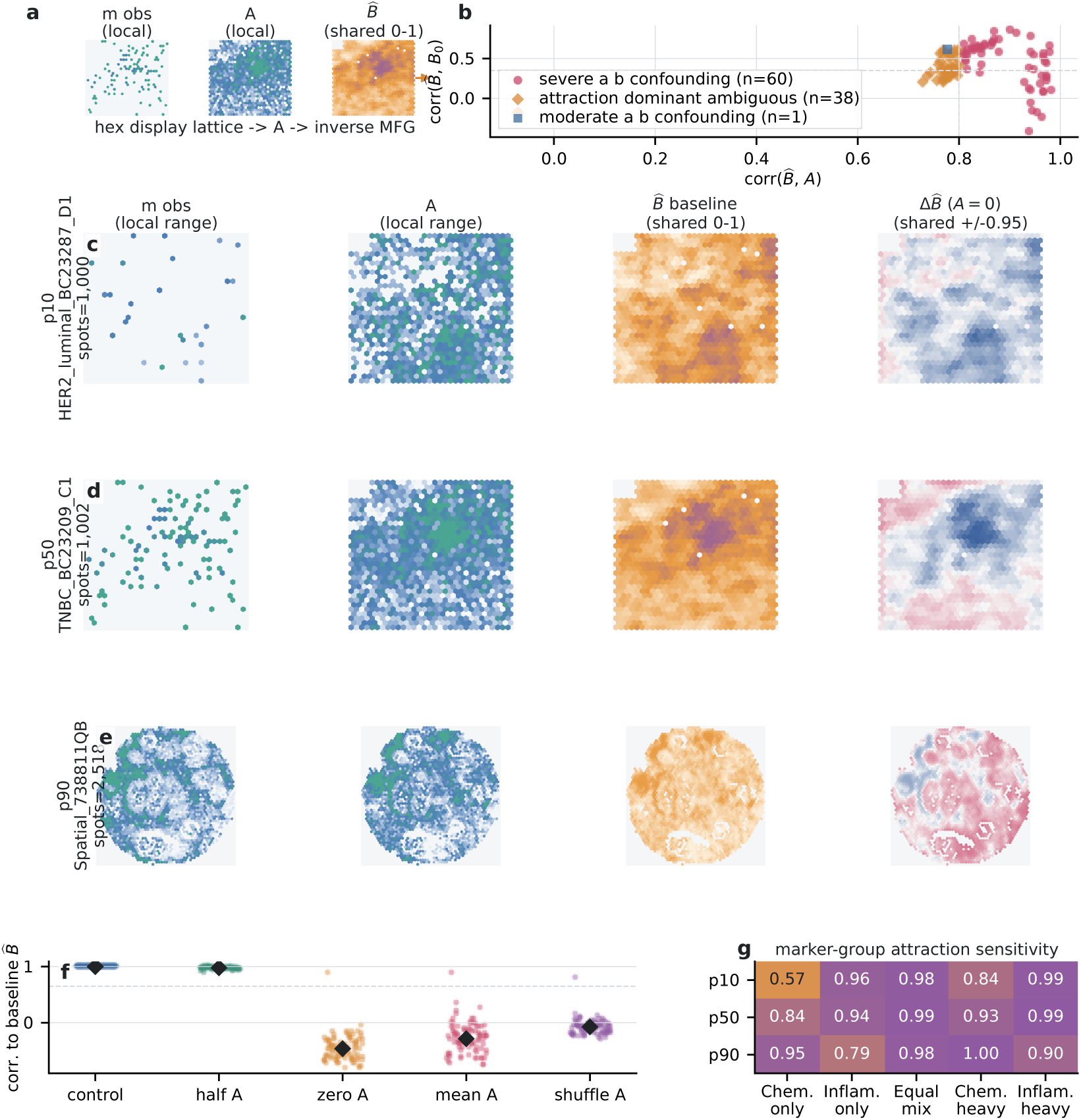
Attraction structure limits identifiability of the equivalent resistance field. a, Real-section input plate showing observed density, specified attraction and inferred equivalent field. **b,** Baseline A/B diagnostics for 99 breast sections; colors and marker shapes denote severe A/B confounding (60), attraction-dominant ambiguity (38) and moderate A/B confounding (1) under the predefined descriptive rules. **c–e,** Sections at the p10, p50 and p90 ranks of baseline attraction correlation, showing observed density and attraction on labelled section-local ranges, baseline *B* on a shared 0–1 range and the *A* = 0 field difference on a shared symmetric *−*0.95 to 0.95 range. Maps in a and c–e use the index-preserving hexagonal display lattice. **f,** Attraction perturbations across all 99 sections, with individual points and medians. **g,**Targeted marker-component sensitivity for the same three sections. Source data and display scales are provided.

The inverse field was then recomputed under four predefined attraction perturbations while holding the observed density, mask and solver contract fixed. Halving attraction amplitude preserved the field relatively well (median field correlation 0.977; median top-20% support IoU 0.786), whereas zero, constant-mean and spatially shuffled attraction produced median field correlations of *−*0.462, *−*0.285 and *−*0.074, respectively (Fig. 4c–e). The complete 99-section distributions show that the spatial structure of attraction, rather than amplitude alone, materially influences the recovered field.

Sections nearest the p10, p50 and p90 ranks of baseline corr(*B, A*) (ranks 11, 50 and 89) were selected for spatial inspection. Real-section panels use the same index-preserving hexagonal display lattice, with missing source indices left blank. Observed-density and attraction maps use labelled section-local ranges; baseline *B* uses a shared 0–1 range, and the *A* = 0 difference maps use a shared symmetric *−*0.95 to 0.95 range (Fig. 4c–e). A targeted marker-component analysis on these sections showed mixed stable, shifted and sensitive outcomes for chemokine-only, inflammation/antigen-only and weighted component definitions (Fig. 4g); Fig. 4f shows all broad perturbation results. Together, these perturbations bound the identifiability of *B* under the current attraction specification and motivate comparison with matched anatomical measurements.

### 2.6 Projection fidelity defines the current applicability boundary

We quantified projection fidelity and forward solvability for all 99 primary breast sections. Mask round-trip intersection-over-union (IoU) measured agreement between source support and its projection onto the shared solver grid. Ninety-six sections produced stabilized re-forward fields, while three failed at the forward-prior stage and remained in the cohort accounting (Fig. 5a–c). Projection IoU did not predict section-level density agreement across the successful sections.

**Figure 5:**
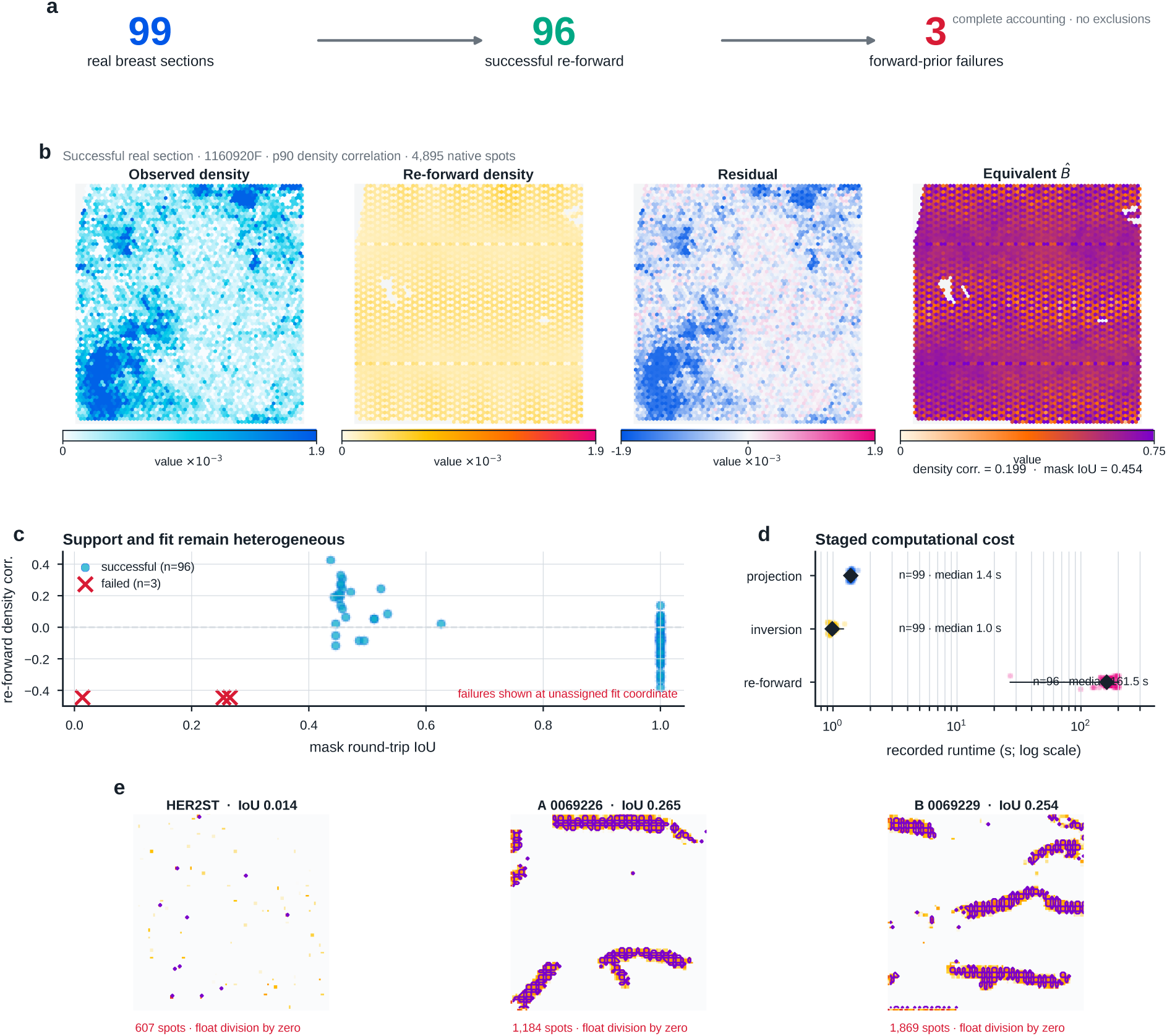
Projection fidelity defines the current applicability boundary. a, Accounting of 99 breast sections: 96 yield stabilized re-forward outputs and three fail during the forward-prior stage. **b,** The p90 successful section, showing observed density, re-forward density, pointwise signed residual and equivalent resistance *B̂* at all 4,895 source spot IDs. Stored array indices are mapped to the index-preserving hexagonal display lattice, with missing indices left blank. Observed and re-forward densities share one range and the residual uses a zero-centred symmetric range. **c,** Re-forward density correlation versus mask round-trip IoU for all 96 successful sections; failures are shown at an unassigned fit coordinate. **d,** Projection (*n* = 99), inversion (*n* = 99) and reforward (*n* = 96) runtime distributions on a logarithmic axis, with medians, interquartile ranges and observed ranges. **e,** Projected mask-probability fields for the three failures, with IoU, spot count and solver reason. Spatial section is the observational unit. Source data are provided.

Projection, inversion and re-forward runtimes are shown with the projected mask-probability fields for the three forward-prior failures (Fig. 5d,e). These results identify projection geometry and forward-prior solvability as measurable limits of the current workflow. Generalization beyond the analysed cohort remains to be evaluated in a held-out breast dataset.

### 2.7 Stabilization, sensitivity and efficiency expose numerical trade-offs

We consolidated the numerical controls that determine whether the inverse workflow is solver-ready. In the paired degraded-observation comparison, stabilization increased the barrier-field correlation from 0.440 to 0.992 and changed the re-forward status from failure to success (Fig. 6a). The complete 21-row ordered ablation contains seven failed re-forward conditions and identifies densityfloor smoothing, with optional value-field smoothing, as the first successful components for the predefined low-noise, high-noise and sparse-spot cases (Fig. 6b). These deterministic cases isolate numerical stabilization effects.

**Figure 6:**
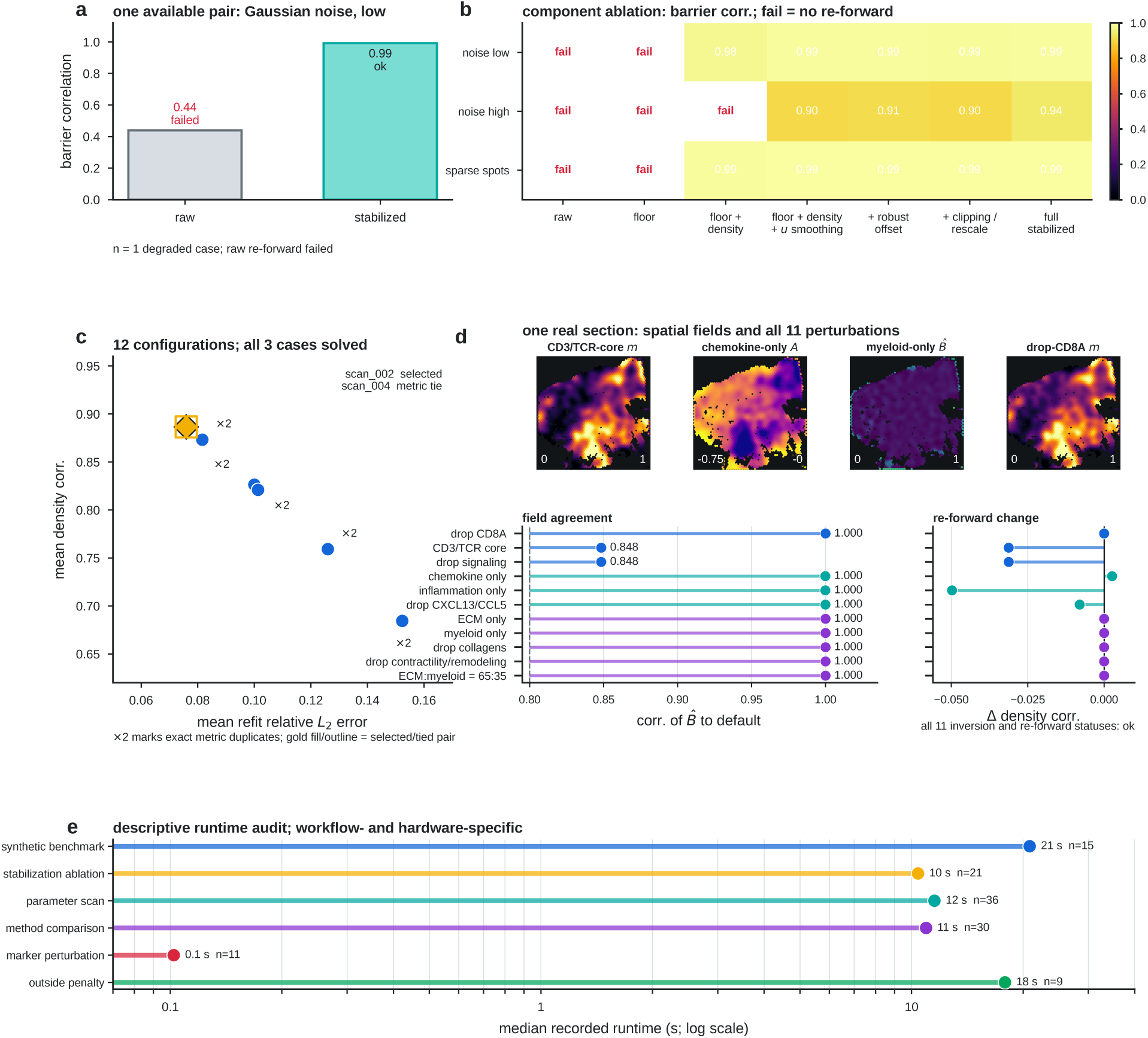
Stabilization, sensitivity and efficiency expose numerical trade-offs. a, Paired raw and stabilized results for the gaussian-noise-low degraded observation. **b,** Complete 21-row ordered stabilization ablation across low-noise, high-noise and sparse-spot synthetic cases; cells show barrier correlation and ‘fail’ denotes unavailable re-forward output. **c,** Twelve-configuration stabilization scan plotted as mean density correlation versus mean refit relative *L*_2_ error; scan002 is selected by the recorded ranking rule and scan_004 is its metric tie. **d,** Four spatial fields sampled at all 3,058 source spot IDs and 11 targeted marker/component perturbations for one real section. Maps use the index-preserving hexagonal display lattice; quantitative panels report inferred-*B* agreement with the default and change in re-forward density correlation. **e,** Median recorded runtime for six workflows on a logarithmic time axis. Panels a–c are deterministic numerical cases, panel d is a single-section sensitivity analysis and panel e reports implementation-specific runtime. Source data are provided.

The 12-configuration stabilization scan exposes a geometry–refit trade-off: the selected scan (scan 002) has mean density correlation 0.887 and mean refit relative *L*_2_ error 0.076, whereas higher barrier correlation occurs with a modestly worse density refit (Fig. 6c). In one real section, an 11-variant marker perturbation shows that inferred-field similarity to the default depends on the observation or context component; the corresponding maps are sampled at all 3,058 source spot IDs and shown on the same hexagonal display lattice (Fig. 6d). Recorded runtimes provide implementation-specific measurements for the six workflows (Fig. 6e).

### 2.8 Complementary native-graph analysis across heterogeneous tumour records

To examine computational portability beyond the primary breast-section PDE workflow, we performed a separate complementary native-graph analysis on 30 retained spatial records representing 29 distinct native bundles. These records were drawn from public spatial datasets spanning breast, colorectal, head-and-neck, ovarian, lung, pancreatic, prostate and intestinal tumour contexts [12–24].

Extended Data Fig. 1 summarizes cohort coverage and screening evidence. Extended Data Figs. 2–4 document input pairing, native-bundle structure and tumour-proxy boundary summaries. Extended Data Figs. 5–8 assess marker-proxy associations, residual terms, parameter sensitivity and perturbation stability. These analyses retain native sampling coordinates and use a fixed zero-current graph operator; they do not use the regular PDE-grid projection or run real-section PDE re-forward calculations. Results are summarized at the record or bundle level and are intended as exploratory computational-portability and sensitivity analyses, rather than a pooled validation of the 99-section breast cohort or an independent biological validation.

### 2.9 Equivalent barrier fields define functional immune-blocking states

Under the model, the inferred barrier fields describe functional immune-blocking states. High-barrier configurations showed concentrated resistance near tumor margins, stronger depletion across peri-tumoral layers and lower predicted access to inner tumor regions. Low-barrier configurations showed weaker resistance fields and broader immune propagation toward malignant tissue. These configurations define a model-based continuum from relatively permissive to strongly blocked states. The equivalent barrier field provides a common representation for the modeled effects of extracellular matrix density, fibrosis, stromal geometry and suppressive signaling on T-cell advance. It summarizes their joint association with immune access under the specified observation, attraction and resistance-prior fields.

The barrier field is therefore interpreted as a model-derived functional state variable. Its correspondence with extracellular matrix, fibrosis, suppressive markers and progression-related endpoints requires evaluation against matched biological measurements.

### 2.10 Noninvasive tumor progression assessment from T-cell distribution-derived tomography

The tissue-derived tomography state could provide a target for future noninvasive assessment. Imaging features, circulating immune markers, liquid-biopsy signals or limited-sampling spatial assays could be mapped to the same state if matched measurements become available.

Such an application would require paired tissue and external measurements, a prespecified prediction task and validation in held-out patients. The present tissue-only analysis defines the candidate target representation but does not test cross-modal prediction.

The immediate output is therefore an MFG-constrained inverse state derived from spatial tissue measurements. Whether that state can support repeated, less-invasive assessment is a testable question for future matched multimodal studies.

## 3 Discussion

The controlled benchmarks establish numerical recoverability under matched model assumptions. Across five single-peak cases, reference–recovery correlations were 0.9962–0.9990, and stabilization restored re-forward solvability in degraded observations. Performance on the heterogeneous breast cohort was substantially weaker: among 96 successful sections, the median observed–re-forward density correlation was *−*0.060. The method therefore recovered controlled fields accurately but did not reproduce immune-density variation consistently across the real cohort under the current parameterization.

Attraction sensitivity identifies a central source of this discrepancy and limits interpretation of the inverse field. Across 99 breast sections, the median correlation of B̂ with the specified attraction field was 0.825, and removing, averaging or shuffling attraction markedly changed the recovered field. Thus, *B̂* currently reflects the joint observation–attraction specification rather than a uniquely identified resistance component. Constraining attraction independently or measuring it from matched data is necessary to separate these contributions.

The numerical controls also define practical limits of the workflow. Density and value-field smoothing improved solvability in the degraded synthetic cases, but three breast sections still failed at the forward-prior stage. Mask round-trip IoU quantified projection fidelity yet did not predict section-level re-forward agreement. These findings separate solver stability and geometric support from biological fit: improving one does not by itself resolve model misspecification in heterogeneous tissue.

Within these limits, the inferred field is useful as an auditable functional representation of modeled resistance to T-cell redistribution. It places boundary accumulation, layered depletion and core access in a common inverse problem and exposes how the result changes with stabilization, projection and attraction specification. Correspondence between this state and extracellular matrix, fibrosis, suppressive signaling or pathological progression remains to be tested using matched measurements.

The current evidence is cross-sectional and section-level. Complete patient grouping was unavailable for several external records, no held-out breast cohort was used to assess generalization and no noninvasive modality was paired with the tissue-derived state. Longitudinal and clinical interpretations therefore require patient-resolved sampling, independently constrained biological fields, matched multimodal measurements and prospective evaluation.

## 4 Figure Legends

### 4.1 Figure 1 — Breast spatial data and the MFG inverse workflow

**a,** Original HT397B1 tissue without an overlay, the same registered tissue with measured immune-module spots, and the corresponding observation, attraction and resistance-prior fields. Constructed fields are oriented to the registered upper-origin view. **b,** Equivalent resistance, solved value field, re-forward density and signed residual on the same section, including calculated local*−∇u* directions; solver fields use a neutral background without tissue-image compositing. **c,** Original tissue images without overlays and the observed, inferred and re-forward outputs for breast and colorectal sections processed through the same deterministic interface; each hexagonal display is oriented to its registered upper-origin pixel axes. **d,** All finite in-mask observed versus re-forward density samples, used to quantify same-section numerical consistency.

### 4.2 Figure 2 — Stabilization preserves controlled resistance geometry and rescues degraded re-forward solves

**a,** Reference resistance fields for five predefined standardized similar-scene cases, with fixed field-quantile contours. **b,** Recovered equivalent fields with sampled lattice points. **c,** Signed re-forward residuals and per-case stored metrics. **d,** High-noise reference, observed, raw and stabilized fields. **e,** Complete 3-by-7 ordered-component success lattice. **f,** Runtime medians and ranges across the deterministic numerical cases.

### 4.3 Figure 3 — Breast-section re-forward fit is heterogeneous across sections and data sources

**a–c,** Direct source-spot observed, sampled re-forward, pointwise residual and sampled equivalent-field maps for the p10, p50 and p90 successful breast sections. Stored array indices are mapped to the index-preserving hexagonal display lattice. Density, residual and equivalent-field families use shared cross-section ranges. **d,** Ranked density correlations for all 96 successful sections, with the three forward-prior failures shown as red crosses. **e,** Source-stratified section-level distributions. Spatial section is the statistical unit, and source comparisons are descriptive.

### 4.4 Figure 4 — Attraction structure limits identifiability of the equivalent resistance field

**a,** Real-section input plate with observed density, specified attraction and inferred equivalent field. **b,** Baseline A/B identifiability diagnostics for all 99 breast sections, with 60 severe confounding, 38 attraction-dominant ambiguous and one moderate confounding label under predefined descriptive rules. **c–e,** Baseline and perturbed fields for p10, p50 and p90 sections on the index-preserving hexagonal display lattice. Observed-density and attraction maps use section-local ranges; baseline *B̂* and the *A* = 0 differences use shared 0–1 and symmetric *−*0.95 to 0.95 ranges, respectively. **f,** Broad attraction perturbation results and medians across all sections. **g,** Targeted marker-component attraction sensitivity. These analyses quantify dependence of *B* on the attraction specification.

### 4.5 Figure 5 — Projection fidelity defines the current applicability boundary

**a,** Complete 99-section accounting, comprising 96 stabilized re-forward results and three forward-prior failures. **b,** Observed, re-forward, residual and equivalent-resistance fields for the p90 successful section at all 4,895 source spot IDs, shown on the index-preserving hexagonal display lattice. **c,** Re-forward density correlation versus mask round-trip IoU for 96 successful sections, with failures shown at an unassigned fit coordinate. **d,** Projection, inversion and re-forward runtime distributions on a logarithmic axis. **e,** Projected mask-probability fields for the three failures, with IoU, spot count and solver reason. The projection metric quantifies numerical support fidelity, and spatial section is the observational unit.

### 4.6 Figure 6 — Numerical controls, sensitivity and efficiency

**a,** Paired raw and stabilized results for the gaussian-noise-low degraded observation. **b,** Complete 21-row ordered stabilization ablation across three deterministic degraded synthetic cases; ‘fail’ denotes unavailable re-forward output. **c,** Twelve-configuration stabilization scan and predefined scan 002 selection, with scan 004 retained as a metric tie. **d,** Four spatial fields sampled at all 3,058 source spot IDs and 11 targeted marker/component perturbations for one real section; maps use the index-preserving hexagonal display lattice, and quantitative panels report inferred-*B* agreement with the default and change in re-forward density correlation. **e,** Median recorded runtimes for six workflows on a logarithmic axis. Panels a–c are deterministic numerical cases, panel d is a single-section sensitivity analysis and panel e reports implementation-specific runtime.

## 5 Methods

### 5.1 Spatial field construction and coordinate handling

For each spatial-transcriptomics section, gene counts were normalized to 10,000 counts per spot and transformed with log(1 + *x*). The immune observation was the mean normalized expression of available T-cell and T-cell-receptor markers (CD3D, CD3E, CD3G, CD2, CD247, CD8A, CD8B, TRAC, TRBC1, TRBC2, LCK and LAT). The attraction signal combined chemokine/recruitment and inflammation/antigen-presentation modules with weights 0.65 and 0.35, and the resistance prior combined extracellular-matrix/stromal and suppressive-myeloid modules with the same weights. An epithelial reference module used EPCAM, KRT8, KRT18, KRT19, MUC1, ERBB2 and MKI67. Marker availability was recorded separately for every section. Each module score was rescaled within the in-tissue spots using its 5th and 95th percentiles and clipped to [0, 1].

The source arrays retain the spot barcode, in-tissue flag, array row and column, and full-resolution pixel coordinates. Formal solver inputs use array row and column as an index grid; pixel coordinates are retained as auxiliary fields. The index grid is projected to the regular PDE grid for inversion and sampled back at the same source spot IDs for display and section-level comparison. Thus, the hexagonal display lattice preserves source-index adjacency, while the PDE solve is performed on the regular projected grid.

### 5.2 Public data sources and cohort construction

All biological inputs were obtained from open spatial-transcriptomics records; no new human specimens were collected for this study. The primary cohort was assembled from the versioned cohort manifest supplied in the accompanying analysis archive before the cohort-level inversion. It contains 99 marker-complete human breast spatial sections from 15 public source records: eight 10x Genomics Visium records (Block A sections 1 and 2, fresh-frozen breast cancer, CytAssist fresh-frozen breast cancer, FFPE ductal carcinoma in situ/invasive carcinoma, whole-transcriptome breast cancer, fluorescent-CD3 invasive ductal carcinoma and CytAssist FFPE gene-and-protein expression) [25–32]; three Figshare records (the Ståhl breast-cancer H5AD collection, HER2ST in the SECTOR benchmark and a breast-cancer metabolic spatial dataset) [33–35]; and four Zenodo records (the breast-cancer atlas, the Andersson HER2-positive benchmark, breast-cancer liver metastasis and the TNBC Visium archive) [36–39].

The 68 Ståhl sections contain 23 verified patient identifiers; patient-level inference was not attempted for the remaining sources because their downloaded metadata do not provide complete patient grouping. Accordingly, the statistical unit throughout the cohort audit is the spatial section. A predeclared sensitivity subset contains 84 sections with at least 1,000 in-tissue spots and 2,000 detected genes. The source records, accessions, sample identifiers, platform descriptions and processing status are retained in the accompanying versioned cohort manifest and primary-cohort table.

The expression matrices, spatial coordinates, tissue images and source annotations were downloaded from the cited records and converted to a common field-bundle interface. For the D2 section from TNBC patient BC23288, the H5AD source points to the upstream Mendeley record containing the paired H&E image and registered spot pixels used in Fig. 1c [40]. The derived fields are analysis representations: m obs is the marker-defined immune observation, attraction is a specified context field, barrier prior is a marker-derived prior and tumor reference is an epithelial reference field. The inverse solver combines these inputs to produce *B*, a model-derived equivalent functional resistance field. The controlled resistance fields and degraded observations used in the numerical controls are generated inputs.

The public records retain complementary biological contexts from their source studies. Ståhl et al. established sequencing-based spatial transcriptomics and supplied the breast sections used in the H5AD collection [6]. Andersson et al. analysed HER2-positive breast cancer with spatial deconvolution and reported tumour-associated cell-type interactions, providing the context for the HER2ST/HER2-positive records [8]. Wu et al. assembled a single-cell and spatially resolved atlas of human breast cancers, which underlies the breast-atlas records in this cohort [9]. Doan et al. developed a cross-modality AI framework for breast-cancer metabolic dysregulation and released the associated spatial data used here [41]. The TNBC archive is associated with the integrative spatial-omics study by Zhu et al., which identified tumour-promoting multicellular niches and immunosuppressive mechanisms across Black American and White American patients [10]. These citations describe the biological provenance and original study context; the persistent Figshare and Zenodo records remain the authoritative download sources for the exact files analysed here. The HER2ST/SECTOR and liver-metastasis records are cited at the repository level because the deposited metadata do not unambiguously identify an additional primary article. Likewise, the 10x Genomics demonstration records are cited as provider datasets rather than assigned unrelated platform papers.

### 5.3 Observation channels for T-cell redistribution

The analysis was organized around three observation channels designed to capture progression-relevant immune access. Boundary infiltration quantified T-cell density within a predefined distance from the tumor margin. Layered infiltration quantified T-cell density across successive peri-tumoral zones extending outward or inward from the boundary. Core penetration quantified T-cell density or occupancy within inner tumor regions.

Joint analysis preserves the complementary information encoded by the three channels. A sample with high boundary infiltration may still be strongly blocked if layered infiltration decays rapidly and core penetration remains low. Conversely, moderate boundary accumulation may indicate relative accessibility if the layered signal persists and the core remains populated. This multichannel description provided the observation target for inverse inference.

### 5.4 Mean-field game model

Let Ω *⊂* R^2^ denote the tissue domain and let *ρ*(*x, t*) denote T-cell density at position *x* and pseudo-time *t*. We represent microenvironmental resistance by an equivalent barrier field *B*(*x*) and optional known tissue-context terms *C*_tissue_(*x*). A representative T cell follows a controlled diffusion process in which movement is influenced by migration effort, density-dependent interaction and local resistance, consistent with the standard mean-field game formulation.

The forward model is expressed as a coupled mean-field game system for the value function *u*(*x, t*) and density field *ρ*(*x, t*),

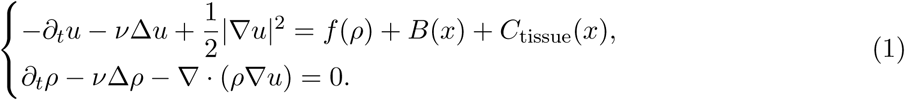

Here *f* (*ρ*) captures interaction or crowding effects and *ν* is the diffusion coefficient. Given a barrier field, the forward system predicts the redistribution of T cells across the boundary region, peritumoral layers and tumor core. The forward dynamics therefore define the mechanistic map from latent resistance structure to observable multichannel T-cell pattern. The finite-element and fixedpoint treatment uses standard numerical discretisations for stationary and time-dependent meanfield games.

### 5.5 Inverse tomographic inference

The inverse problem seeks the barrier field and associated parameters whose forward solution best reproduces the observed multichannel redistribution pattern. We formulate this as a PDE-constrained optimization problem,

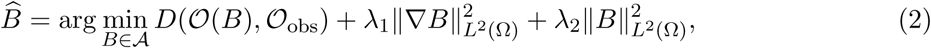

where *O*_obs_ contains the measured boundary, layered and core signals and *O*(*B*) is the corresponding model prediction. The regularization terms control smoothness and amplitude of the inferred barrier field. Derived outputs include spatial barrier maps, boundary exclusion strength, layered blocking strength and core-penetration-related state variables. These outputs define the tomographic state reported in the Results.

### 5.6 Numerical forward and inverse experiments

Numerical experiments were used to test the forward map, inverse recovery and tissue-shaped end-to-end consistency of the MFG framework. Unless otherwise stated, experiments used finite-element resolution nx=18, sampling grid grid n=72, five continuation steps and a maximum of 20 fixed-point iterations. Common model parameters were diffusion coefficient *ν* = 0.04, crowding coefficient *β* = 0.25, barrier strength *λ*_barrier_ = 1.4 and total mass equal to 1.0.

Forward simulations first evaluated whether controlled barrier geometries shaped immune-density equilibria. Five synthetic tumor configurations were considered: simple core, multifocal barrier, irregular lobulated barrier, twin-core bridge and peripheral arc. A separate slice-like forward experiment used a real tissue-derived proxy field and compared the simulated immune density with the observed T-cell proxy under baseline, higher-diffusion, stronger-crowding, weaker-barrier and stronger-barrier parameter settings.

Inverse experiments evaluated whether immune-density observations were sufficient to recover latent barrier structure. Five related single-peak barrier fields were used to generate forward immune-density observations, after which the MFG-constrained inverse solver recovered the barrier field and a re-forward density was computed. Recovery was summarized by relative barrier-field error, barrier correlation, relative re-forward density error and pairwise distances between recovered barriers. A final tissue-shaped proxy experiment tested the full forward–inverse–re-forward chain under a real slice-derived geometry.

### 5.7 Complementary native-graph analysis

The complementary native-graph analysis was performed separately from the primary breast-section PDE workflow. It retained native sampling coordinates, used a fixed zero-current graph operator and summarized results at the record or bundle level. It did not use the regular PDE-grid projection, run real-section PDE re-forward calculations or pool the 30 retained records with the 99-section breast cohort. Tumour and microenvironment quantities in this analysis are post hoc marker proxies and are not treated as inverse inputs or orthogonal biological validation.

### 5.8 Baseline and ablation analyses

The reported numerical controls and sensitivity analyses assessed the behaviour of the inferred representation beyond the default workflow, including stabilization, attraction-component and marker-component perturbations. These analyses focused on barrier recovery, re-forward consistency and dependence on the specified observation and attraction fields. Separate density-only, MFG-removal, barrier-prior-removal, layered-channel and resampling-based validation experiments were not performed.

### 5.9 Association with progression-related states

Potential associations between barrier fields and tissue-level features of immune blocking and progression were treated as exploratory targets. The present study did not evaluate pathological stage, clinical outcomes or matched biological measurements of extracellular matrix, fibrosis or suppressive signalling.

### 5.10 Noninvasive assessment framework

The tissue-derived tomographic state was treated as a target representation for future lower-invasive assessment. In this framework, tissue data would provide supervision for mapping external measurements to progression-related barrier states. Candidate external inputs include imaging features, circulating immune markers, liquid biopsy features or limited-sampling spatial assays. A future cross-modal model would estimate the tissue-derived state from these less-invasive signals.

### 5.11 Statistics and reproducibility

The reported analyses used section-level or record-level summaries appropriate to each workflow. Robustness was assessed through the reported sensitivity, stabilization, attraction-perturbation, projection and native-graph analyses. Sample sizes, medians, correlations and failure counts are reported with the relevant Results and figure legends; biological-group comparisons, uncertainty intervals and resampling-based validation were not part of this section-level study.

## 6 Data Availability

All biological data analysed in this study are publicly available. The primary cohort comprises 99 marker-complete human breast spatial sections from 15 public records, accessed between 6 and 9 July 2026. The 10x Genomics records are Block A sections 1 and 2, fresh-frozen breast cancer, CytAssist fresh-frozen breast cancer, FFPE ductal carcinoma in situ/invasive carcinoma, whole-transcriptome breast cancer, fluorescent-CD3 invasive ductal carcinoma and CytAssist FFPE gene-and-protein expression [25–32]. The Figshare records are the Ståhl breast-cancer H5AD collection (accession 19768351), the HER2ST/SECTOR processed dataset (accession 32029830) and the breast-cancer metabolic spatial dataset (accession 22337722) [33–35]. The Zenodo records are the breast-cancer atlas (record 4739739), the Andersson HER2-positive benchmark (record 15024747), breast-cancer liver metastasis (record 18306028) and the TNBC 10x Visium archive (record 15252874) [36–39]. The cited bibliography entries provide the persistent landing page or DOI for each source record; access conditions and licences remain those of the original providers.

The accompanying analysis archive identifies every dataset ID, sample ID, accession, platform, section count and quality-control field. The complementary native-graph analysis used 30 retained spatial records representing 29 distinct native bundles from 22 public source datasets. Corresponding record mappings, processed field bundles, marker-audit tables, projection summaries, re-forward summaries and figure-level source-data tables are included in the analysis archive. The raw expression matrices and tissue images can be retrieved from the cited accessions using the documented data-preparation materials. The synthetic resistance fields and degraded observations used for numerical controls are generated from the stated configurations. The archived *B* fields are model-derived equivalent functional MFG resistance representations.

## 7 Code Availability

The modelling, inverse-inference, cohort-construction and figure-generation code is included in the accompanying versioned repository and source-data archive. Processing notes and versioned manifests document the conversion from each public source record to the field-bundle interface. A persistent repository DOI for the final code and derived source-data archive will be added to the published version.

---

**Extended Data Fig. 1.**
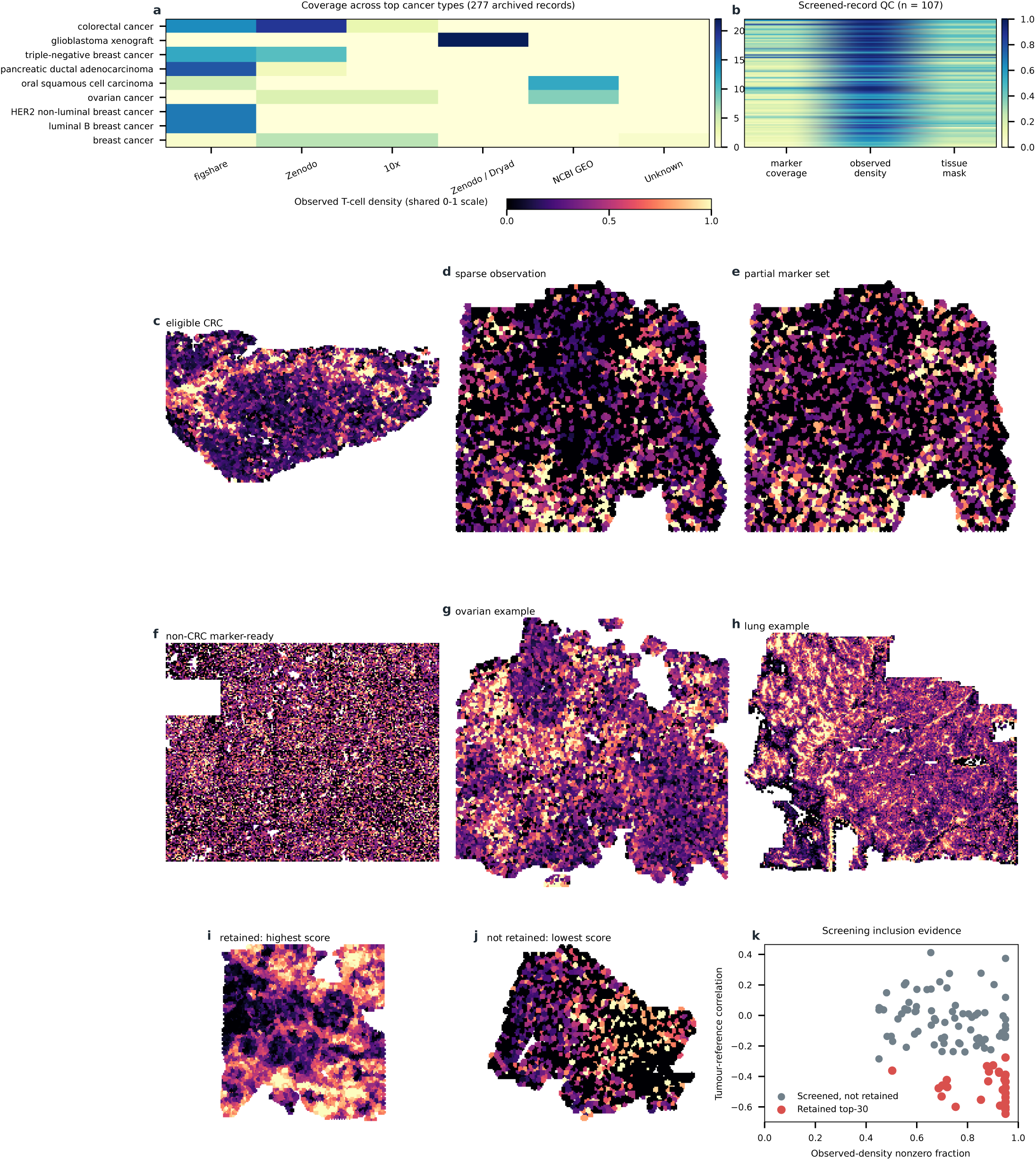
Cohort coverage, screening evidence and observed-density fields. a, Coverage across 277 source records. b, Recorded quality-control fractions for the 107 screened records. c-h, Six deterministic screening roles: eligible colorectal, sparse observation, partial marker set, marker-ready non-colorectal, ovarian and lung. The paired ovarian assays in d-e are distinct assays and are not independent tissues. i-j, Highest-score retained and lowest-score screened-not-retained contrasts; screening non-retention does not imply biological failure. k, Inclusion evidence. Bounded nearest-point cells are a constant-value display support, not capture footprints; missing observations are not filled. The rank-6 CRC map uses its source index-coordinate grid; all other display coordinates are source pixel coordinates. The retained analysis set contains 30 spatial data records.

**Extended Data Fig. 2.**
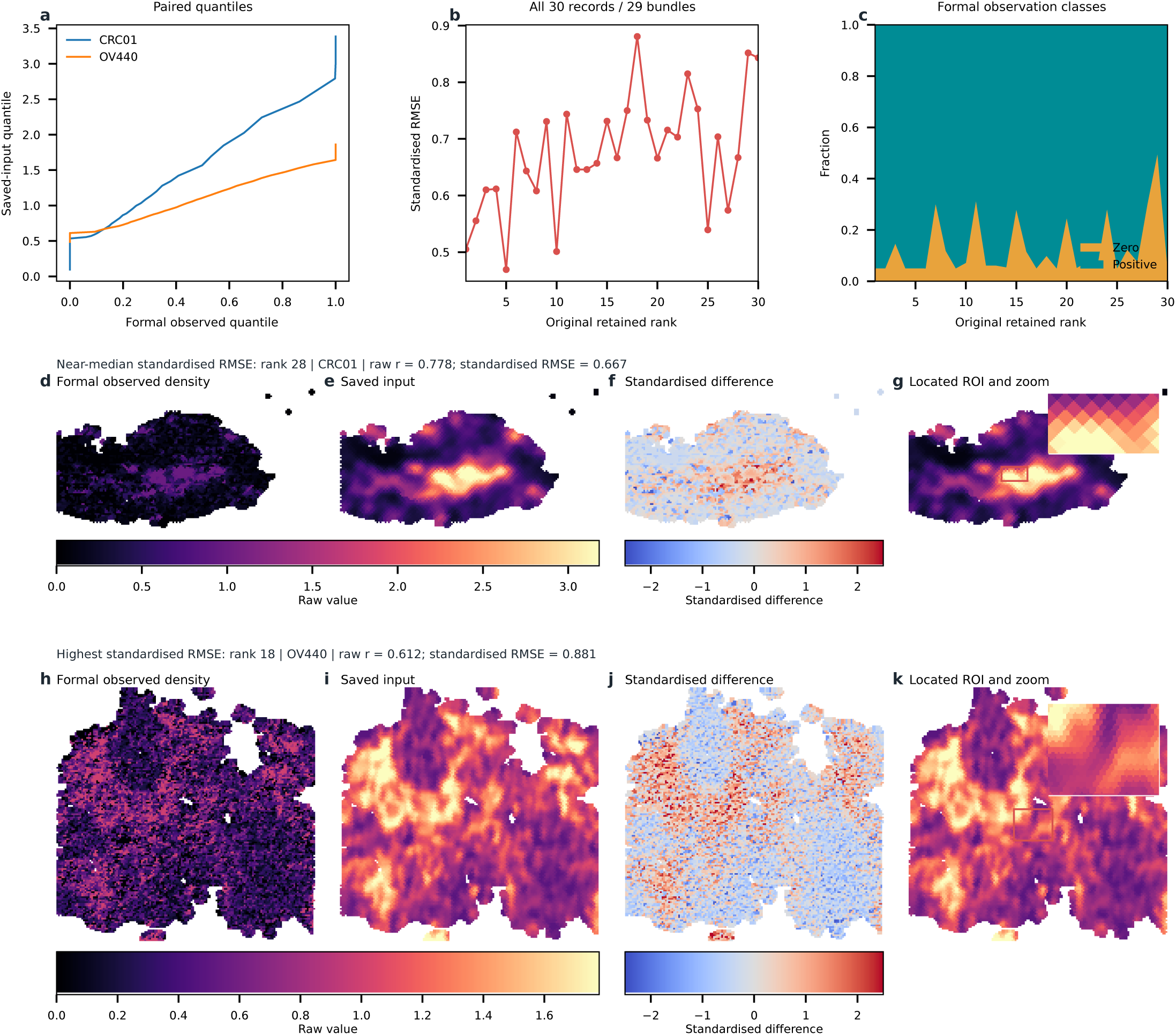
Exact formal-observation to analysis-input pairing. a, Paired raw-observation and analysis-input quantiles for the two displayed records. b, Standardised RMSE across 30 retained records. c, Formal zero and positive observation fractions by retained rank. d-g, Near-median-RMSE CRC01, whose formal computation used a source index-coordinate grid. h-k, Highest-RMSE OV440, with validated source-pixel-matched geometry. Each spatial field uses bounded nearest-point cells with constant source values and no interpolation. Formal observations are re-ordered to analysis coordinates through the validated zero-distance, one-to-one mapping; mapping, raw-value and coordinate hashes are supplied in Source Data.

**Extended Data Fig. 3.**
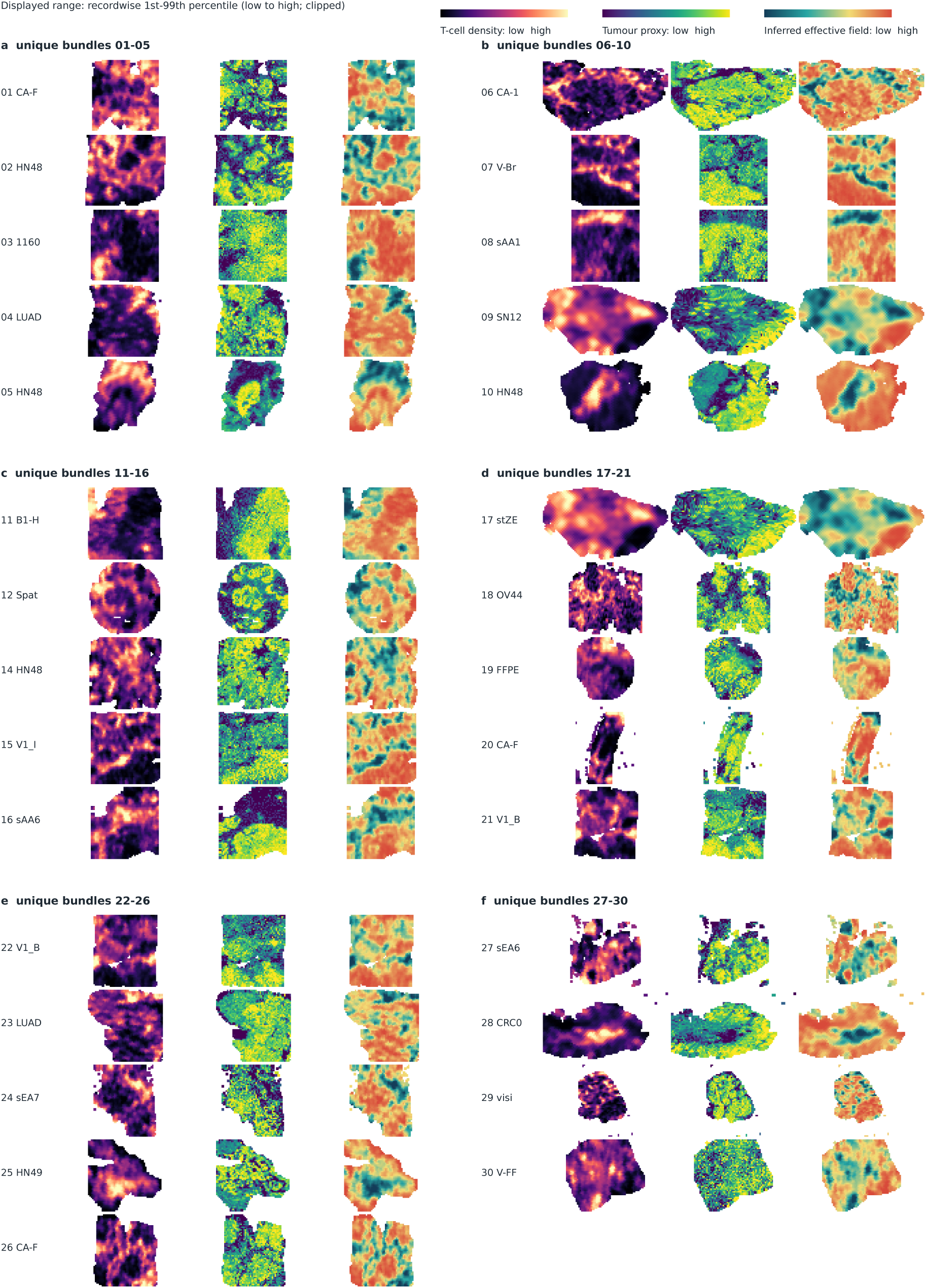
Native-coordinate atlas of the unique retained bundles. The retained analysis set contains 30 record names representing 29 distinct native bundles; ranks 12 and 13 refer to the same bundle. The atlas displays the 29 unique bundles once in original screening order; rank 13 is omitted from the spatial plate but remains in the source-data files and analysis accounting. Each row shows observed normalised T-cell density, tumour proxy, and recomputed inferred effective field. Each observed location is displayed as a bounded nearest-point support cell with radius 0.72 times the record-specific median nearest-neighbour spacing; cells are clipped by native-coordinate neighbour boundaries. This display introduces no interpolation, smoothing or missing-value fill. Full identifiers, source accessions, display endpoints and renderer parameters are provided in the source-data files. Colours span the within-record 1st–99th percentile and are not intended for between-record comparison.

**Extended Data Fig. 4.**
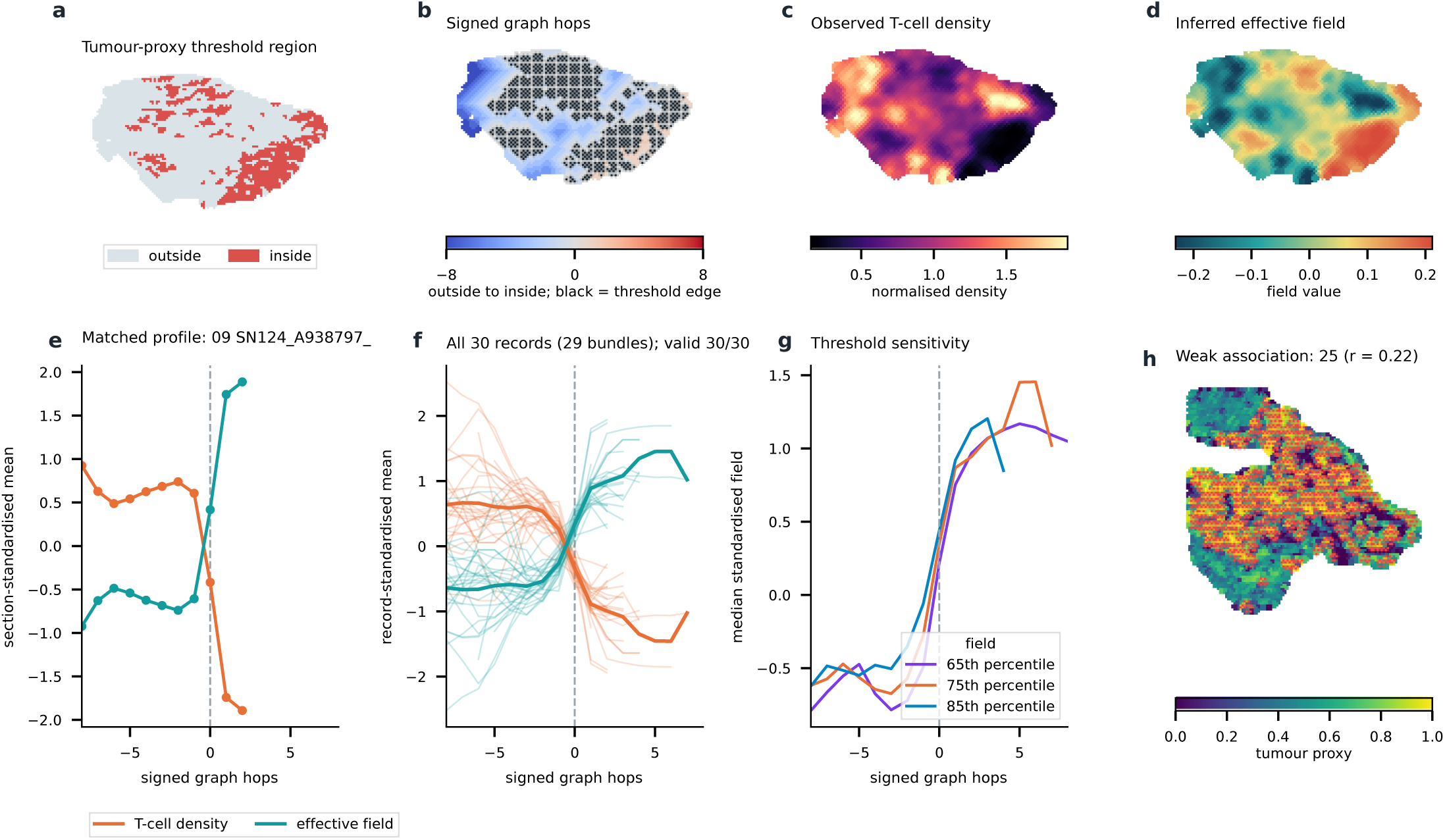
Graph-derived relation to a tumour-proxy threshold boundary. A representative record displays the within-record 75th-percentile tumour-proxy region, graph boundary and signed hops, observed density, and inferred effective field. Spatial values are shown as bounded nearest-point supports with radius 0.72 times the median nearest-neighbour distance; this display does not interpolate values and the supports are not real cell boundaries. Matched profiles include all 30 ordered records. Boundary and distance layers are post hoc graph summaries, not pathology truth. Missing interfaces remain NA.

**Extended Data Fig. 5.**
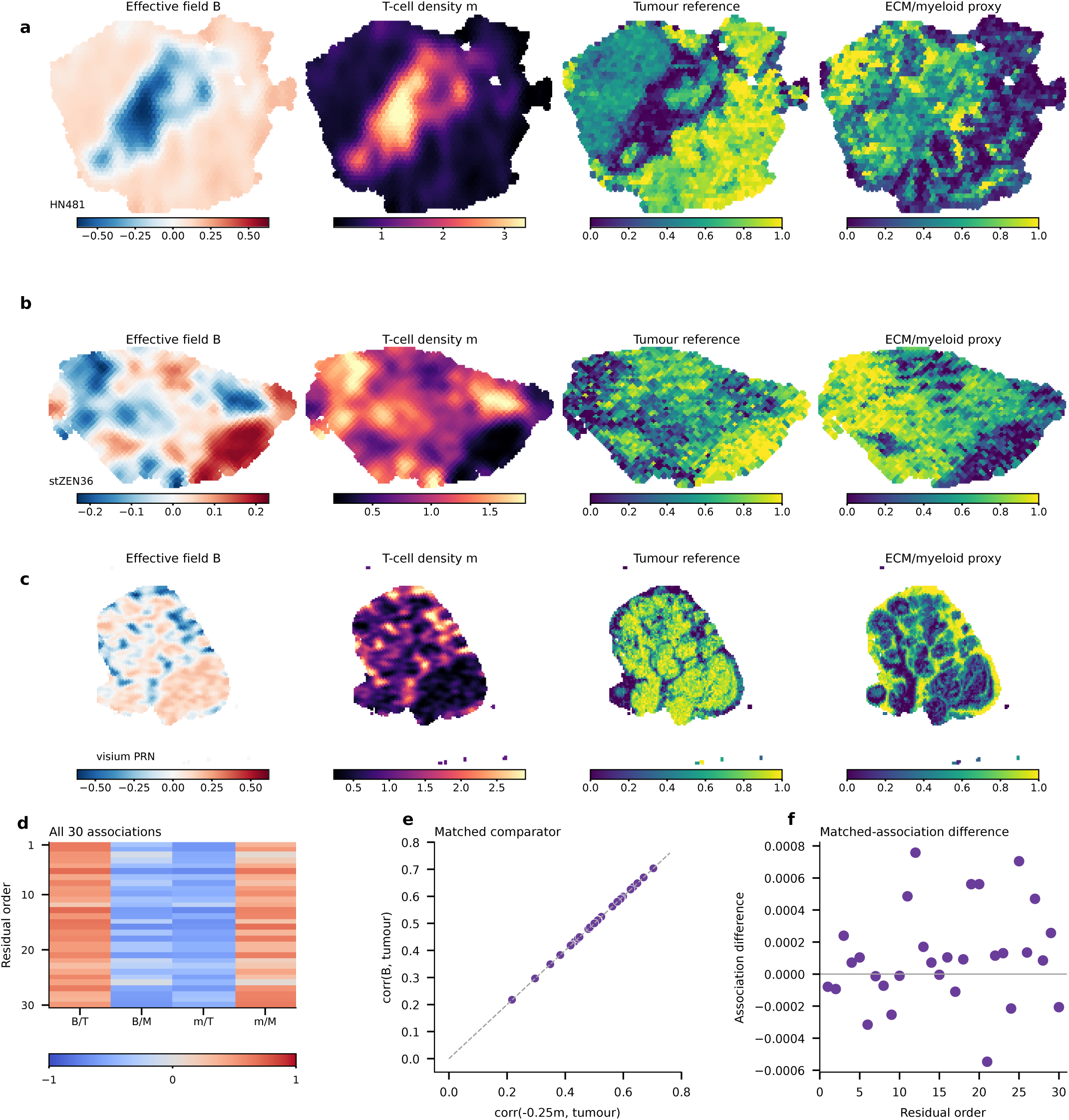
Recomputed inverse fields and spot-aligned microenvironment references. a-c, Low-, median-and high-residual records show effective field B, processed T-cell density, tumour reference and the formal-bundle composite microenvironment score. The latter is a 0.65 ECM/stromal and 0.35 suppressive-myeloid marker-score composite, sampled only where native spot coordinates matched exactly. d, All-record correlations with tumour and microenvironment references. e, Matched density-baseline versus B tumour-reference associations. f, Difference between the paired associations, ordered by direct residual fraction. Colourbars show displayed numeric ranges for every map. Every spatial display uses a constant-valued 32-sided support centred at each native spot: its initial radius is 0.72 times the median native nearest-neighbour spacing and it is clipped at neighbouring perpendicular bisectors. Geometry is built for every native spot before non-finite values are masked; no interpolation, smoothing or gap filling is applied. The analysis comprises 30 spatial data records, and the results are summarized at the record level. These are marker proxies, not orthogonal validation.

**Extended Data Fig. 6.**
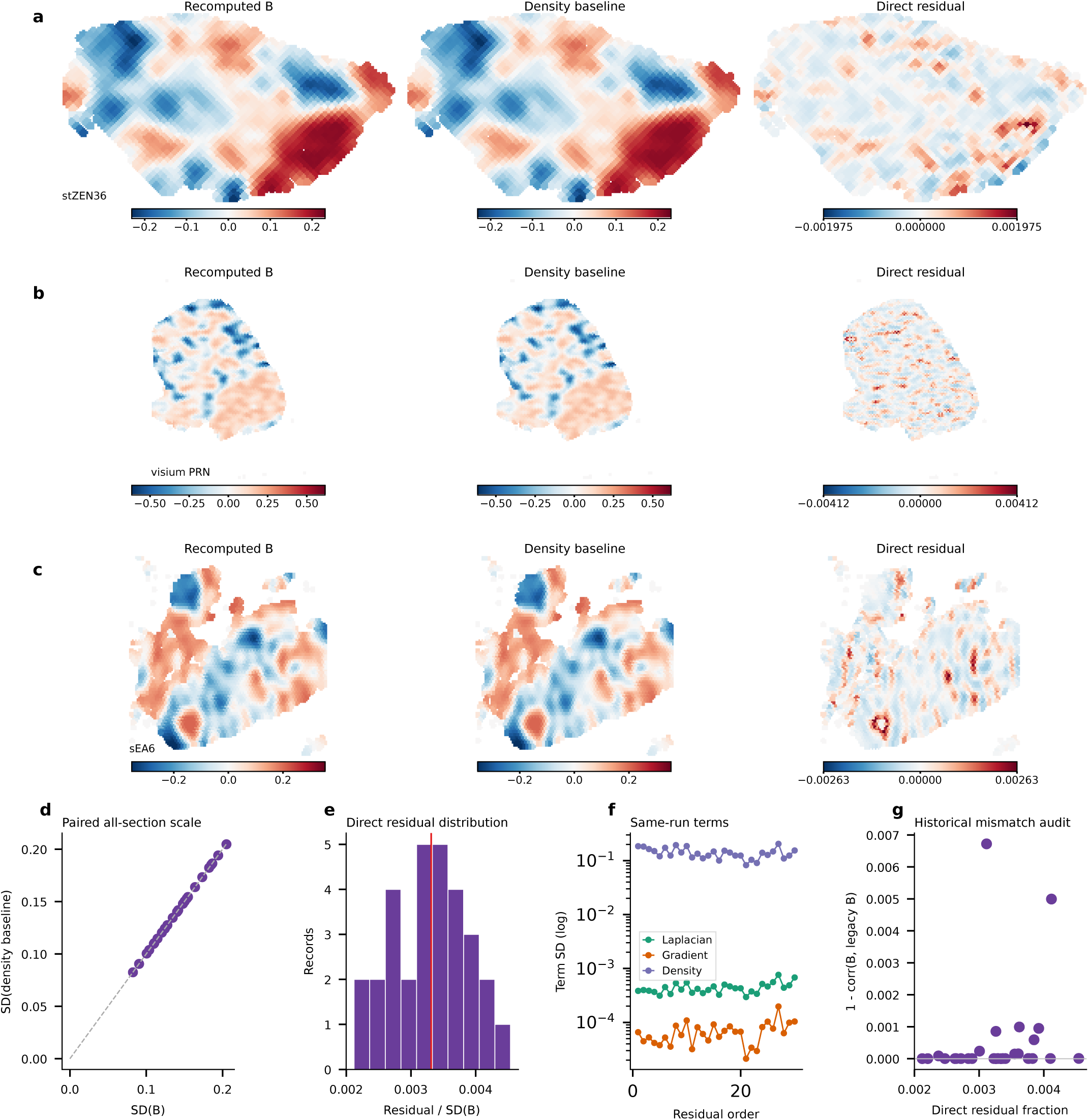
Direct residual audit with same-run inverse terms. a-c, Recomputed B, density baseline and their direct residual for a typical record and the two largest-residual records. Within each record, B and its density baseline use the same zero-centred displayed range; the residual uses a separate symmetric zero-centred range. Numeric colourbars are shown for every map. d, Paired all-record B and baseline scale. e, Direct residual distribution. f, Non-additive term standard deviations plotted separately on a logarithmic scale. g, Reference-field difference as a provenance comparison only. All maps retain native spot coordinates and use constant-valued 32-sided support cells, initialized at 0.72 times median native nearest-neighbour spacing then clipped at neighbouring perpendicular bisectors. Full support geometry is calculated before finite-value masking, so missing values remain missing regions; no interpolation, smoothing or gap filling is applied. The analysis comprises 30 spatial data records, and the results are summarized at the record level.

**Extended Data Fig. 7.**
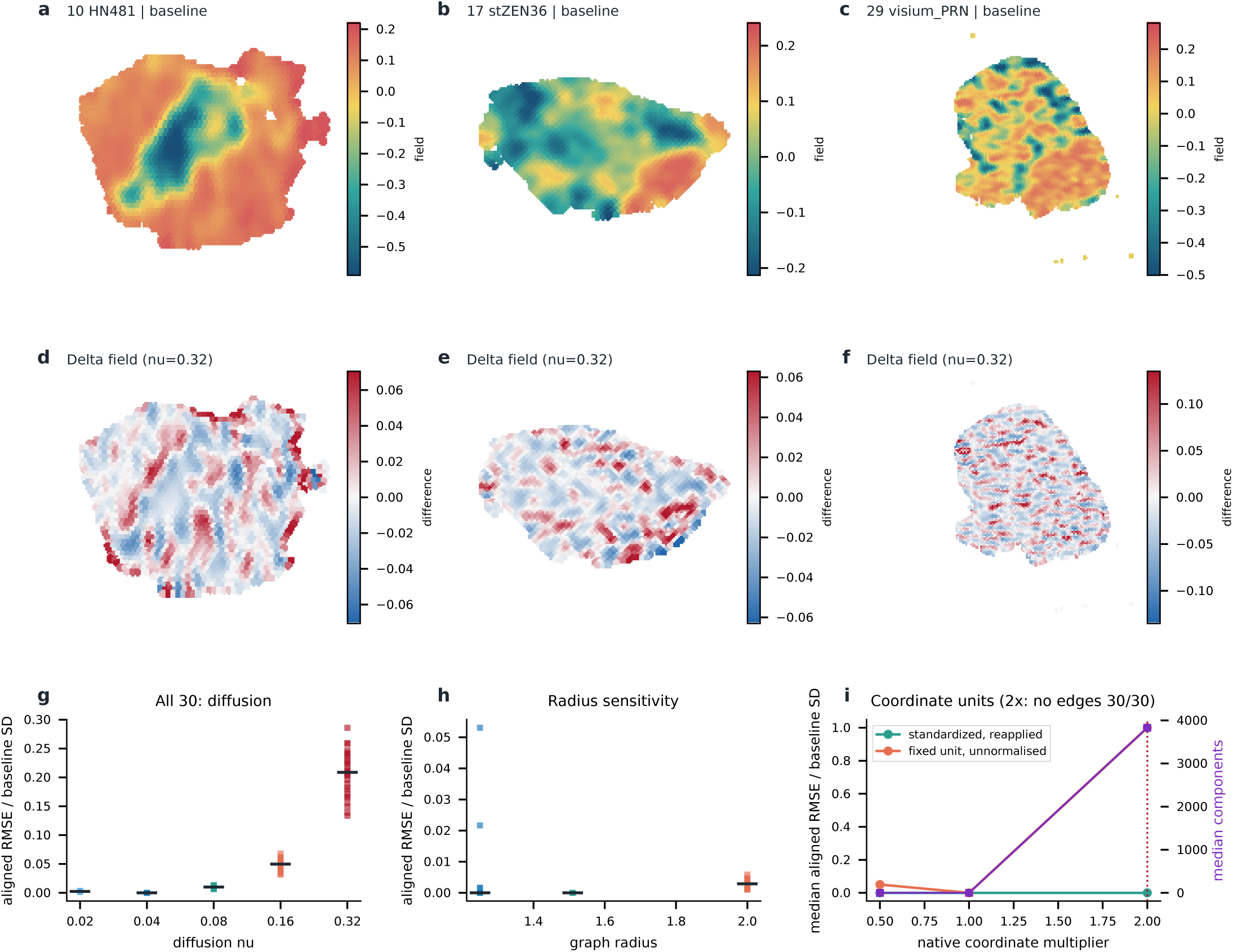
Parameter, topology and coordinate-unit sensitivity of the inferred effective field. Baseline fields and gauge-aligned changes at nu = 0.32 are shown for low, median and high all-record discrepancy examples selected before plotting (a–f). Each difference map has its own symmetric numeric colour scale to preserve local magnitude. Across 30 records, each point is one record and the horizontal mark is the median (g,h). The median component count is one at each displayed radius. Radius changes alter graph connectivity and can alter the inferred field beyond a correlation-only comparison. Coordinate scaling is evaluated in two separate pipelines (i): reapplying section median-neighbour standardisation yields invariance by construction, whereas retaining the baseline length unit changes the graph operator; at 2x all 30 fixed-unit graphs have no edges and are reported as degenerate. Differences remove the mean within each baseline connected component. The analysis is a direct inverse sensitivity analysis under the fixed zero-current formulation; it does not rerun a forward model or interpret the field as a physical barrier. Bounded nearest-point display supports are clipped by neighbour bisectors and limited to a radius of 0.72 times the median nearest-neighbour spacing; they are not biological cell boundaries, and values are not interpolated.

**Extended Data Fig. 8.**
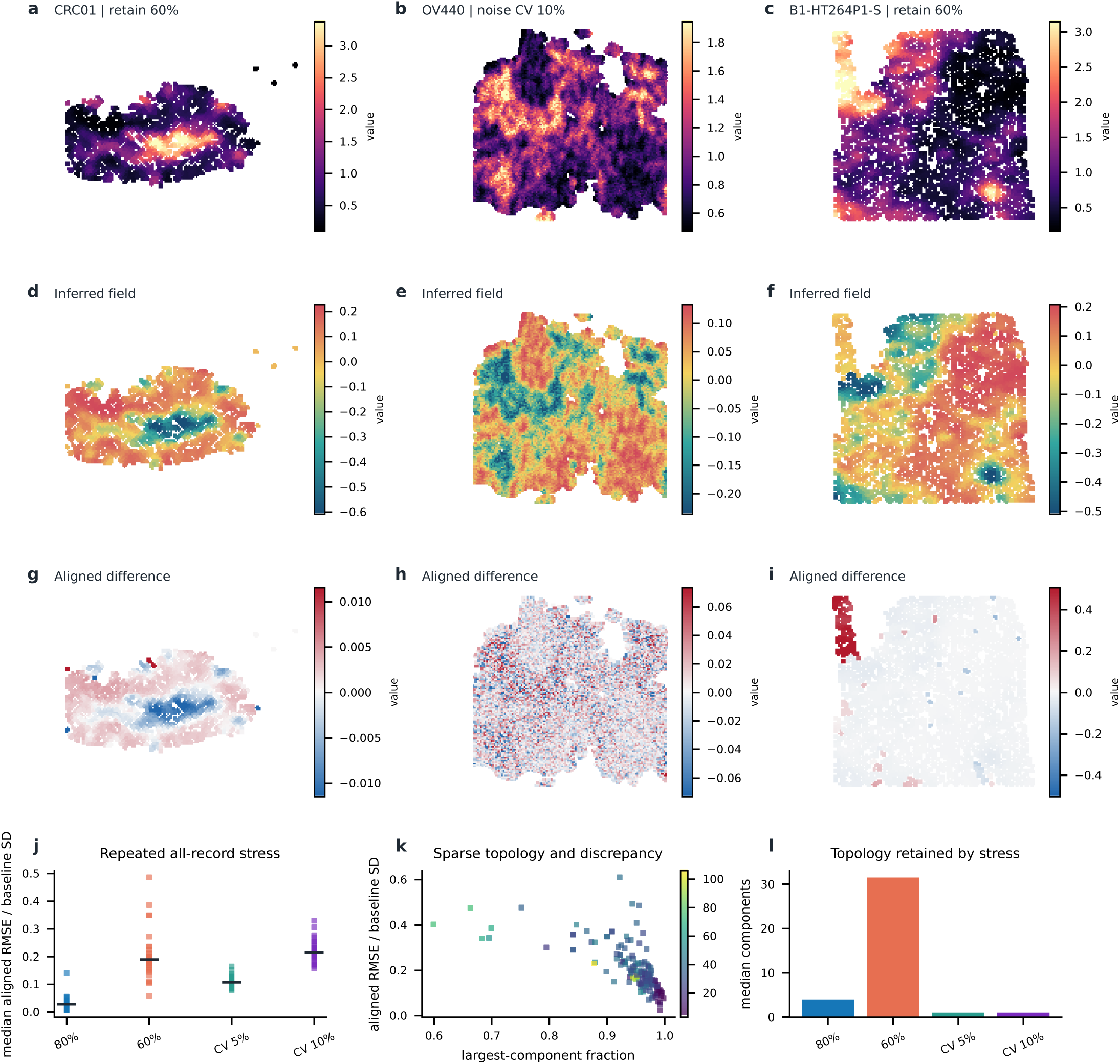
Stability of inferred effective fields under incomplete and noisy observations. Maps show actual retain60% or CV10% perturbed observations, direct inverse fields and common-node differences (a–i). The retain60% columns are selected as the lowest and highest median discrepancies over five fixed seeds; the CV10% column is the highest median CV10% discrepancy. Differences remove offsets within baseline connected components but retain discrepancy from changed candidate-component normalization and topology; they are pipeline discrepancies, not pure gauge-free robustness scores. In j, each point is one record after taking its median over five fixed seeds (30 records per condition). In k, each point is one record-seed result (30 records x 5 fixed seeds = 150 runs); fixed seeds are perturbation realisations, not biological replicates. Panel l reports medians in the displayed condition order: retain80%, retain60%, CV5%, CV10%. Independent random thinning retains original node identities and fixed baseline coordinate units. Multiplicative log-normal perturbations have mean one and CV 5% or 10%; they are stress tests rather than calibrated measurement models. Topology, finite denominators and failure counts are supplied in source data. No forward model is run. Bounded nearest-point display supports are clipped by neighbour bisectors and limited to a radius of 0.72 times the median nearest-neighbour spacing; they are not biological cell boundaries, and values are not interpolated. Supports are constructed on the complete baseline coordinates before thinning; removed observations leave their original supports blank.

